# Discoidin Domain Receptor 2 regulates AT1R expression in Angiotensin II-stimulated cardiac fibroblasts via fibronectin-dependent Integrin-β1 signaling

**DOI:** 10.1101/2021.05.06.442987

**Authors:** Allen Sam Titus, Harikrishnan Venugopal, Mereena George Ushakumary, Mingyi Wang, Randy T. Cowling, Edward G. Lakatta, Shivakumar Kailasam

## Abstract

Recent reports on the cardioprotective effects of fibronectin inhibition following myocardial injury suggest a largely unexplored role for the extracellular matrix (ECM) glycoprotein in cardiac fibroblast function. We probed the molecular regulation and functional implications of fibronectin gene expression in cardiac fibroblasts exposed to Angiotensin II, a major pro-fibrotic factor in the myocardium. Using gene knockdown and over-expression approaches, western blotting and promoter pull-down assay, we show that collagen type I-activated Discoidin Domain Receptor 2 (DDR2) mediates Angiotensin II-stimulated transcriptional up-regulation of fibronectin by Yes-activated Protein in cardiac fibroblasts. Further, siRNA-mediated fibronectin knockdown attenuated Angiotensin II-stimulated expression of collagen type I and anti-apoptotic cIAP2, and enhanced susceptibility to apoptosis. Importantly, an obligate role for fibronectin was observed in Angiotensin II-stimulated expression of AT1R, the Angiotensin II receptor, which would link ECM signaling and Angiotensin II signaling in cardiac fibroblasts. Moreover, conditioned medium from DDR2- or fibronectin-silenced cardiac fibroblasts reduced AT1R expression in H9c2 cardiomyoblasts. The regulatory role of fibronectin in Angiotensin II-stimulated cIAP2, collagen type I and AT1R expression was mediated by Integrin-β1-integrin-linked kinase signaling. In vivo, we observed modestly reduced basal levels of AT1R in DDR2-null mouse myocardium, associated with the previously reported reduction in myocardial Integrin-β1 levels. The role of fibronectin, downstream of DDR2, could be a critical determinant of cardiac fibroblast-mediated wound healing following myocardial injury. In summary, our findings suggest a complex mechanism of regulation of cardiac fibroblast function involving two major extracellular matrix proteins, collagen type I and fibronectin, and their receptors, DDR2 and Integrin-β1.

## Introduction

Cardiac fibroblasts are importantly involved in wound healing and tissue remodelling in the injured myocardium (1). In a setting of myocardial damage, normally quiescent cardiac fibroblasts, under the influence of pro-fibrotic factors such as Angiotensin II (Ang II), are phenotypically transformed into α-smooth muscle actin (α-SMA)-positive myofibroblasts that infiltrate the site of injury, proliferate and produce collagen and other extracellular matrix (ECM) components, leading to the formation of collagenous scar tissue that aids wound repair and preserves myocardial integrity in the short-term. However, unlike in non-cardiac tissues, active myofibroblasts in the heart resist apoptosis and persist in the infarct scar long after termination of the healing response, which results in excessive collagen deposition, adverse tissue fibrosis and compromised ventricular compliance in the long-term (1). Clearly, delineation of molecular mechanisms underlying cardiac fibroblast function and cardiac fibrosis is a clinically relevant goal.

Although earlier studies on cardiac fibrosis had focused on collagen type I as a major contributor to the progression of tissue fibrosis (2), there is now increasing appreciation of the role of cellular fibronectin, an ECM glycoprotein, as an important regulator of ECM and tissue remodelling (3). In fact, recent studies have considered fibronectin as a potential target to limit tissue fibrosis (4). Inhibition of fibronectin polymerization is reported to reduce collagen deposition into the ECM and exert cardioprotective effects through attenuation of cardiac fibroblast proliferation and adverse fibrotic remodelling of the heart in an ischemia-reperfusion model (4). These observations suggest a role for fibronectin in cardiac fibroblast function and cardiac fibrosis and provide compelling rationale for further studies on the regulation of fibronectin expression in cardiac fibroblasts and the mechanisms that underlie its regulatory role in these cells.

We had recently reported that Angiotensin II (Ang II), a potent pro-fibrotic factor whose intracardiac levels are increased following myocardial injury (5), enhances the expression of Discoidin Domain Receptor 2 (DDR2), a collagen receptor tyrosine kinase, which, upon activation by collagen, has an indispensable regulatory role in the expression of collagen type I in Ang II-stimulated cardiac fibroblasts (2, 6). Ang II also induces anti-apoptotic cIAP2 that protects cardiac fibroblasts against ambient stress (7). Interestingly, the pro-survival role of Ang II in cardiac fibroblasts via cIAP2 was also found to be mediated by collagen type I-activated DDR2 (7).

As a sequel to our earlier findings, we now report that DDR2 has an obligate role in Yes-activated Protein (YAP)-mediated transcriptional up-regulation of fibronectin expression by Ang II in cardiac fibroblasts. Downstream of DDR2, fibronectin mediates Ang II-stimulated collagen type I expression and cIAP2-dependent apoptosis resistance in cardiac fibroblasts exposed to oxidative stress. Importantly, fibronectin has an obligate role in Ang II-stimulated expression of the Ang II receptor, AT1R, which would link ECM signaling to Ang II signaling that profoundly impacts cardiac fibroblast function. Further, a combination of gene knockdown and over-expression approaches revealed that the regulatory role of fibronectin in cardiac fibroblasts is mediated by Integrin-β1 signaling. Notably, we observed a small but significant reduction in basal levels of AT1R in DDR2-null mouse myocardium, associated with a reduction in myocardial Integrin-β1 levels reported by us earlier (6). We propose that the pro-survival role of fibronectin and its regulatory role in collagen and AT1R expression, downstream of DDR2, could be critical determinants of cardiac fibroblast response to injury.

## Results

### DDR2 mediates Ang II-stimulated fibronectin expression in cardiac fibroblasts

Treatment of cardiac fibroblasts with Ang II (1 μM) enhanced fibronectin expression (~2-fold), with concomitant increase (~5-fold) in DDR2 expression (Figure 1A). Since enhanced DDR2 expression correlated with enhanced fibronectin expression, the effect of siRNA-mediated DDR2 knockdown on fibronectin expression was checked to ascertain if DDR2 regulates fibronectin. DDR2 knockdown significantly diminished Ang II-induced increase in fibronectin expression (Figure 1B). Further, inhibition of collagen type I-dependent activation of DDR2 using WRG-28 (11) blocked Ang II-stimulated fibronectin expression (Figure 1C). Conversely, over-expression of DDR2 in unstimulated cardiac fibroblasts led to up-regulation of fibronectin expression (Figure 1D), confirming the indispensable role of DDR2 in fibronectin expression.

**Figure 1:**
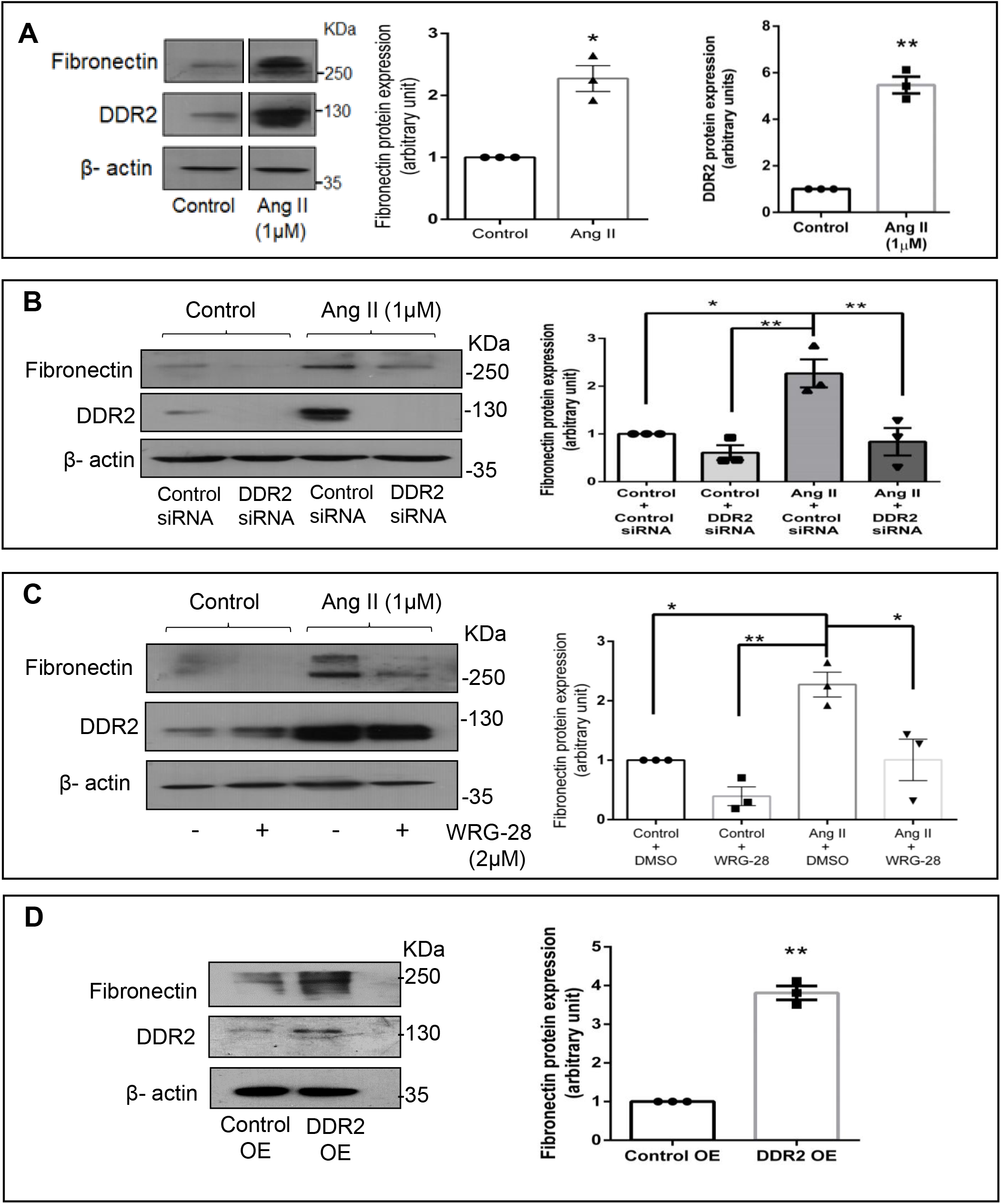
DDR2 mediates Ang II-stimulated fibronectin expression in cardiac fibroblasts. **(A)** Sub-confluent quiescent cultures of cardiac fibroblasts were stimulated with Angiotensin II (Ang II) (1 μM). Protein was isolated at 12 h of Ang II treatment and subjected to western blot analysis for detection of fibronectin and DDR2, with β-actin as loading control. *p< 0.05 and **p< 0.01 vs. control. (**B**) Cardiac fibroblasts were transiently transfected with DDR2 siRNA (5 pmol) or control (scrambled) siRNA prior to treatment with Ang II for 12 h. Fibronectin protein expression was examined, with β-actin as loading control. Validation of DDR2 silencing is also shown. *p<0.05, **p< 0.01 (comparisons as depicted in the Figure). (**C**) Cardiac fibroblasts were treated with WRG-28 (2 μM) for 1 h prior to Ang II treatment. Cells were collected at 12 h post-Ang II treatment and fibronectin protein levels were examined, with β-actin as loading control. *p< 0.05 and **p< 0.01 (comparisons as depicted in the Figure). (**D**) Cardiac fibroblasts were transfected with DDR2 cDNA over-expression plasmid (DDR2 OE) (with empty vector control, Control OE), as described under Methods, and fibronectin protein expression was examined, with β-actin as loading control. **p< 0.01 vs. Control OE. Validation of DDR2 over-expression is also shown. Data are representative of 3 independent experiments, n=3, Mean ± SEM.

### Transcriptional regulation of fibronectin by DDR2-activated YAP transcription co-activator

Since YAP was earlier reported to be activated downstream of DDR2 (6), and bioinformatics analysis of the fibronectin promoter region showed a consensus binding site for TEAD (a YAP-co-activated transcription factor), we checked whether YAP inhibition would have an effect on fibronectin expression. YAP inhibition using a chemical inhibitor (verteporfin) or siRNA reduced Ang II-stimulated fibronectin expression (Figures 2A,B). Further, chromatin immunoprecipitation assay (ChIP) demonstrated YAP binding to the promoter region of fibronectin (containing the TEAD consensus sequence) in Ang II-treated cells, which was attenuated in DDR2-silenced (siRNA) or inhibited (WRG-28) cells (Figure 2C), confirming transcriptional regulation of fibronectin by DDR2-activated YAP.

**Figure 2:**
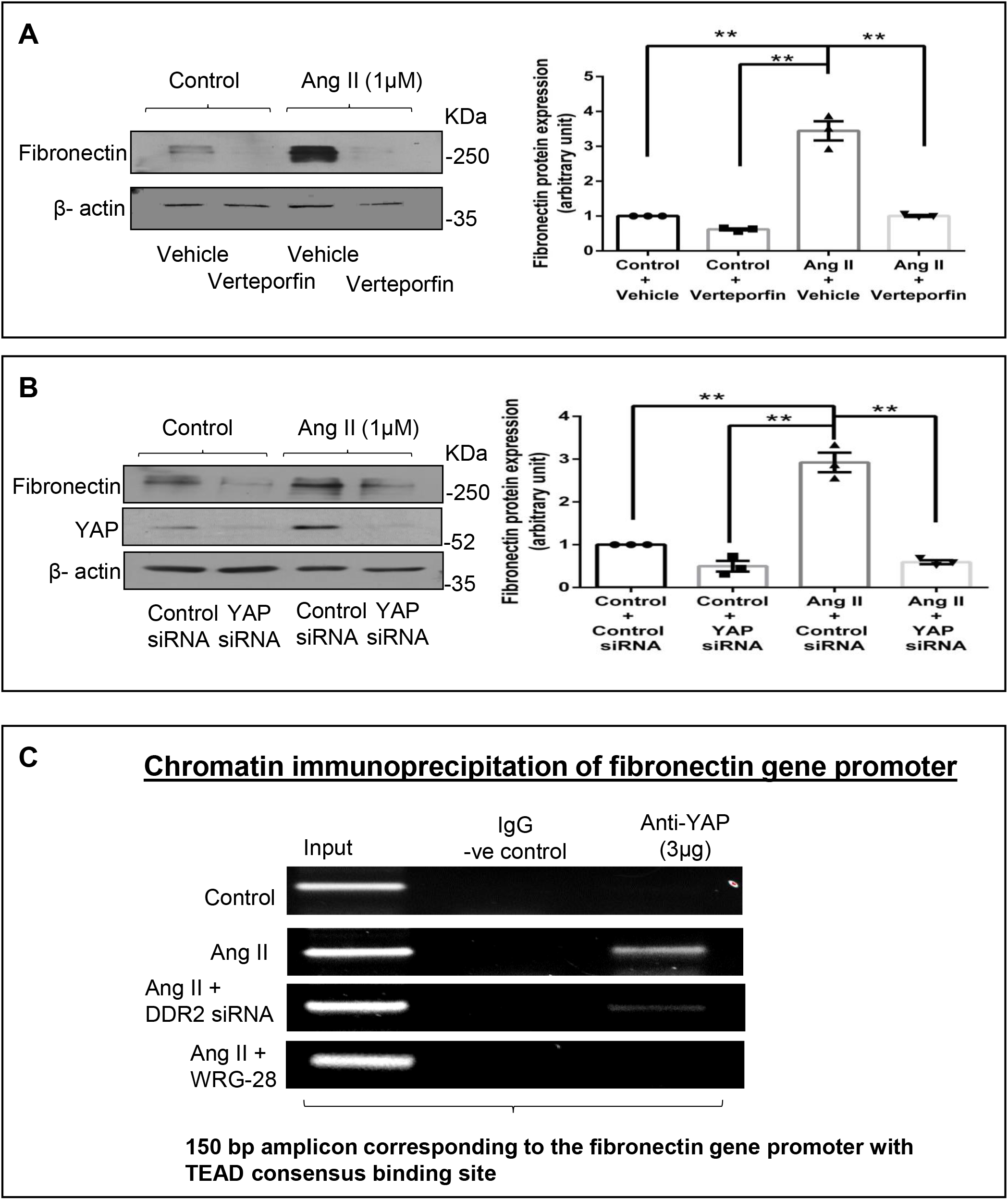
Transcriptional regulation of fibronectin by DDR2-activated YAP transcription coactivator. (**A**) Cardiac fibroblasts were treated with Verteporfin (10 μM) for 1 h prior to Ang II treatment. Cells were collected at 12 h post-Ang II treatment and fibronectin protein levels were examined, with β-actin as loading control. **p<0.01 (comparisons as depicted in the Figure). (**B**) Cardiac fibroblasts were transiently transfected with YAP siRNA (10 pmol) or control (scrambled) siRNA prior to treatment with Ang II for 12 h. Fibronectin protein expression was examined, with β-actin as loading control. Validation of DDR2 silencing is also shown. ** p< 0.01 (comparisons as depicted in the Figure). (**C**) Sub-confluent quiescent cultures of cardiac fibroblasts were transiently transfected with YAP siRNA (10 pmol) or control (scrambled) siRNA prior to treatment with Ang II (1 μM) and another set of cells were pre-treated with WRG-28 for 1 h. Cells were collected at 30 min post-Ang II treatment and chromatin was immunoprecipitated by anti-YAP antibody, followed by PCR amplification, and analysed on a 2% agarose gel for presence of the 150 bp region of fibronectin gene promoter containing the TEAD consensus binding site.

### Functional significance of fibronectin expression in cardiac fibroblasts

#### i) A role for fibronectin in Ang II-dependent cIAP2 expression and apoptosis resistance in cardiac fibroblasts exposed to oxidative stress

Fibronectin has been reported to enhance cell survival in various cell types (12–15). Previous studies from our laboratory had demonstrated that Ang II enhances cIAP2 expression in cardiac fibroblasts that in turn protects these cells against ambient stress (7). In the present study, the role of fibronectin in the regulation of Ang II-dependent cIAP2 expression in cardiac fibroblasts was probed. Fibronectin knockdown using siRNA attenuated Ang II-stimulated cIAP2 expression (Figure 3A) and enhanced apoptosis susceptibility of cardiac fibroblasts exposed to 25 μM H_2_O_2_, as indicated by decreased pro-caspase 3 and increased cleaved-caspase 3 levels (Figure 3B). The observations pointed to the role of fibronectin in enhancing cardiac fibroblast survival under conditions of oxidative stress.

**Figure 3:**
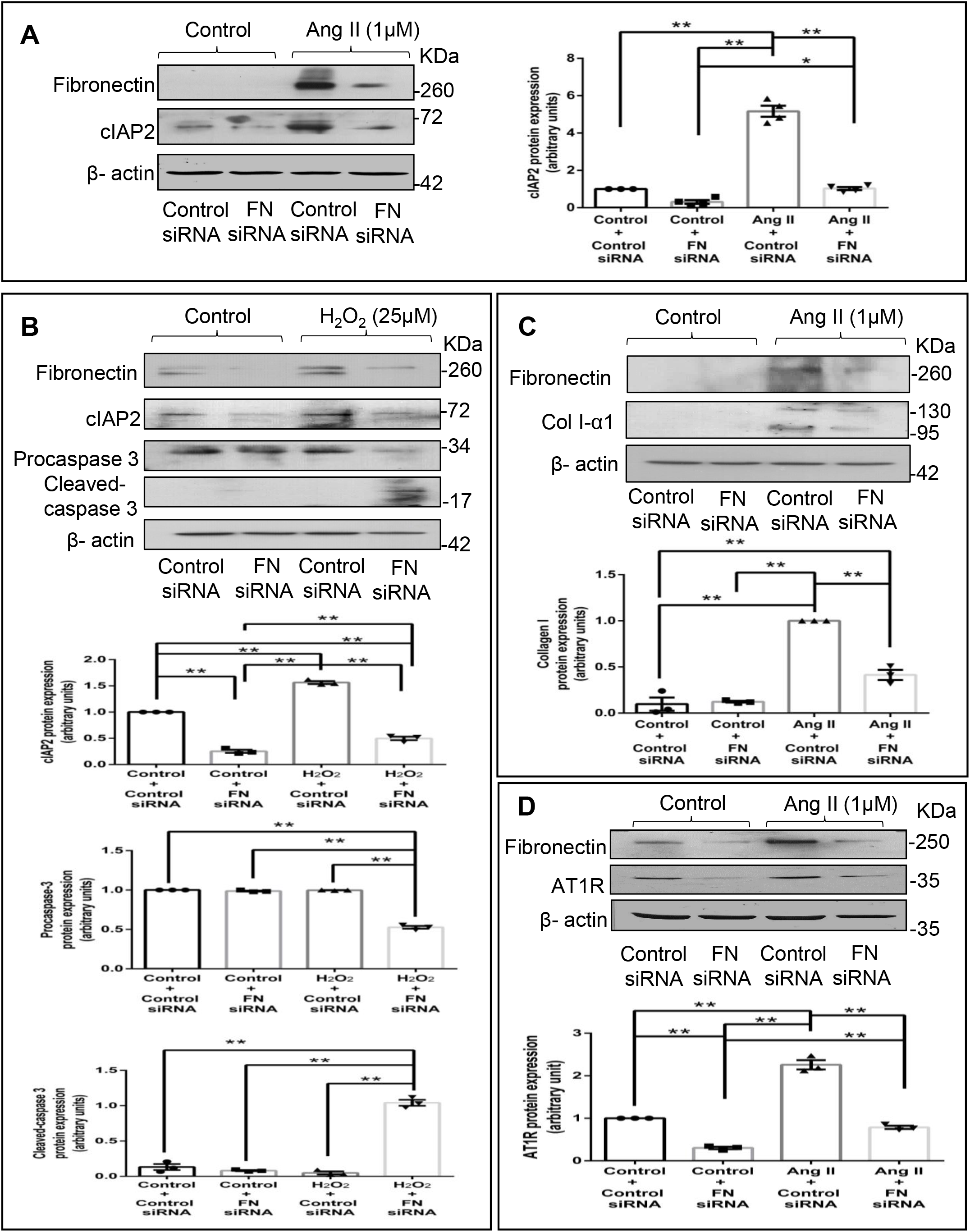
Fibronectin mediates Ang II-stimulated expression of cIAP2, collagen type-I and AT1R in cardiac fibroblasts. (**A**) Cardiac fibroblasts were transiently transfected with fibronectin siRNA (20 pmol) or control (scrambled) siRNA prior to treatment with Ang II for 12 h. cIAP2 protein expression was examined, with β-actin as loading control. Validation of fibronectin silencing is also shown. *p< 0.05 and **p< 0.01 (comparisons as depicted in the Figure). (**B**) Cardiac fibroblasts were transiently transfected with fibronectin siRNA (20 pmol) or control (scrambled) siRNA. Following treatment of cells with 25 μM H_2_O_2_ for 3 h, the cells were collected and levels of procaspase 3 and cleaved-caspase 3 were determined, with β-actin as loading control. Validation of fibronectin silencing and associated changes in cIAP2 levels are also shown. **p< 0.01 (comparisons as depicted in the Figure). (**C,D**) Cardiac fibroblasts were transiently transfected with fibronectin siRNA (20 pmol) or control (scrambled) siRNA prior to treatment with Ang II for 12 h. (**C**) Collagen type 1 protein expression was examined, with β-actin as loading control. Validation of fibronectin silencing is shown. **p< 0.01 (comparisons as depicted in the Figure). (**D**) AT1R protein expression was examined, with β-actin as loading control. Validation of fibronectin silencing is shown. **p< 0.01 (comparisons as depicted in the Figure). Data are representative of 3 independent experiments, n=3. Mean ± SEM.

#### ii) Fibronectin mediates Ang II-dependent increase in collagen type I expression

Fibronectin has been implicated in hepatic and pulmonary fibrosis (16–19), and global or fibroblast-specific ablation of fibronectin in adult mice post-injury resulted in significant cardio-protection with reduced hypertrophy and fibrosis (4, 20). Therefore, the possible role of fibronectin in collagen type I expression in cardiac fibroblasts was probed next. Significant attenuation of Ang II-stimulated collagen type I expression (Figure 3C) upon RNA interference-mediated knockdown of fibronectin suggested a role for fibronectin in the regulation of collagen expression in cardiac fibroblasts.

#### iii) Fibronectin mediates Ang II-stimulated AT1 receptor expression in cardiac fibroblasts

Previous studies from our laboratory had demonstrated that Ang II acts via the AT1 receptor (AT1R) to enhance cIAP2 and collagen type I expression in cardiac fibroblasts (7, 21, 22). Therefore, we checked whether fibronectin plays a role in AT1R expression in Ang II-treated cardiac fibroblasts. Interestingly, fibronectin knockdown down-regulated both basal and Ang II-stimulated AT1R expression (Figure 3D), clearly indicating the role of fibronectin in AT1R expression in cardiac fibroblasts.

### Fibronectin regulates Ang II-mediated AT1R expression through Integrin-β1/Integrin-linked kinase signaling pathway

Fibronectin is reported to act via integrin α5β1 (23), and inhibition of the β1 subunit, the partially-conserved catalytic subunit of integrins (24–26), rather than the α5 subunit, has a more pronounced inhibitory effect on the binding and assembly of exogenous fibronectin (23, 27). In the present study, we observed that fibronectin siRNA attenuates Integrin-β1 expression in Ang II-treated cardiac fibroblasts (Figure 4A), suggesting that fibronectin has an obligate role in the regulation of its receptor. The effect of knockdown of Integrin-β1 and Integrin-linked kinase (ILK), a proximal effector of Integrin-β1 (28), on AT1R was probed next. Abrogation of Integrin-β1 signaling using Integrin-β1 or ILK siRNA led to attenuation of AT1R expression in Ang II-stimulated cardiac fibroblasts (Figure 4B-C). Notably, over-expression of Integrin-β1 in fibronectin-silenced cells partially restored the expression of cIAP2, collagen type I and AT1R in Ang II-treated cells (Figure 4D), clearly demonstrating the involvement of fibronectin-dependent Integrin-β1 signaling in the regulation of cIAP2, collagen type I and AT1R expression in cardiac fibroblasts.

**Figure 4:**
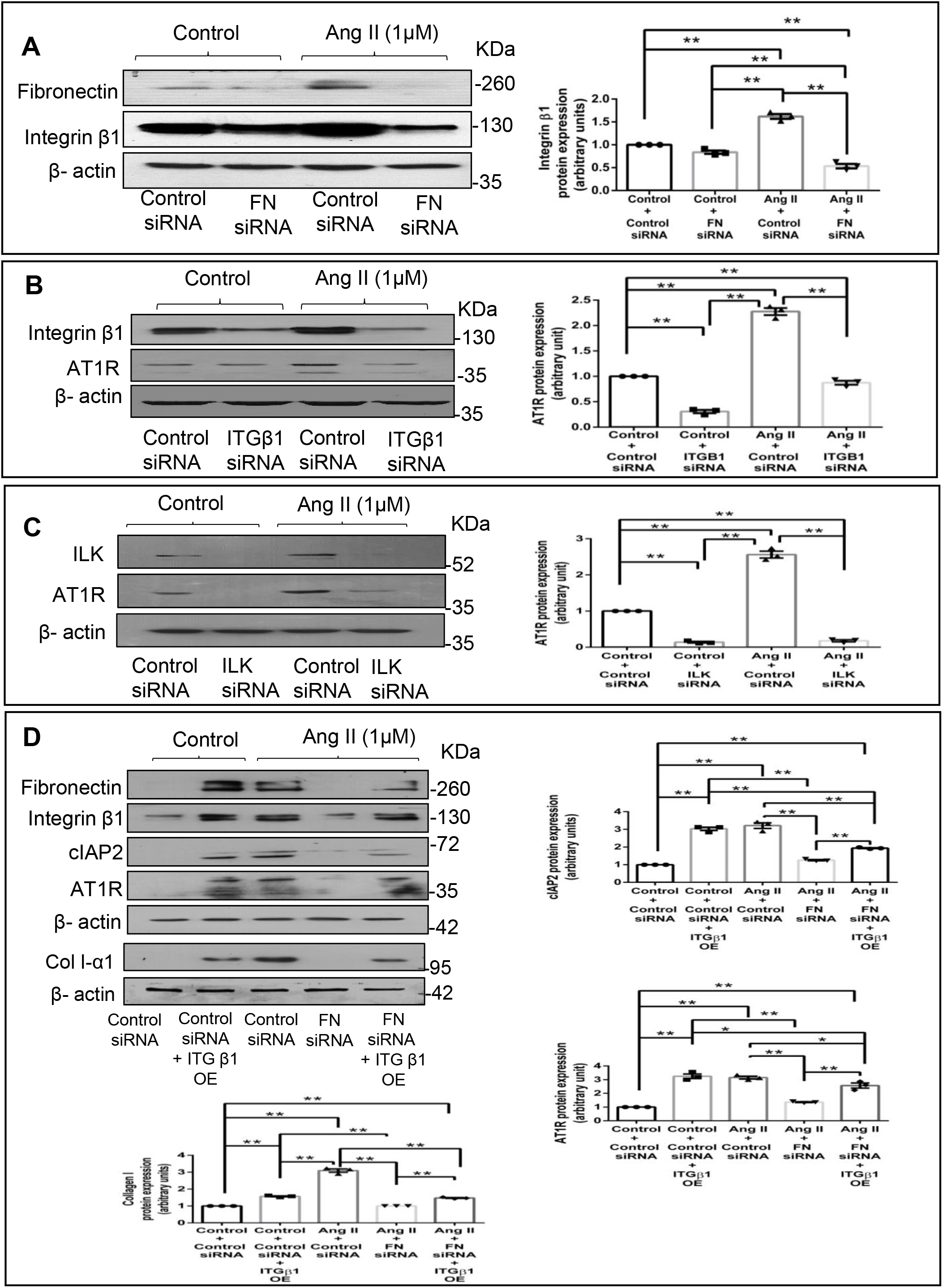
Fibronectin regulates Ang II-stimulated AT1R expression through Integrin β1/Integrin-linked kinase signaling pathway. (**A**) Cardiac fibroblasts were transiently transfected with fibronectin siRNA (20 pmol) or control (scrambled) siRNA prior to treatment with Ang II for 12 h. Integrin-β1 (ITGB1) protein expression was examined, with β-actin as loading control. **p< 0.01 (comparisons as depicted in the Figure). (**B**) Cardiac fibroblasts were transiently transfected with ITGB1 siRNA (5 pmol) or control (scrambled) siRNA prior to treatment with Ang II for 12 h. AT1R protein expression was examined, with β-actin as loading control. Validation of Integrin-β1 silencing is also shown. **p< 0.01 (comparisons as depicted in the Figure). (**C**) Cardiac fibroblasts were transiently transfected with ILK siRNA (5 pmol) or control (scrambled) siRNA prior to treatment with Ang II for 12 h. AT1R protein expression was examined, with β-actin as loading control. **p< 0.01 (comparisons as depicted in the Figure). (**D**) Cardiac fibroblasts were co-transfected with fibronectin siRNA (10 pmol) and ITGB1 cDNA over-expression plasmid (2 μg). Following transfection, the cells were revived in M199 with 10% serum for 12 h. Post-revival, the cells were serum-deprived for 24 h prior to treatment with Ang II for 12 h. Cells were collected and protein levels of AT1R, cIAP2 and collagen type 1 were examined, with β-actin as loading control. *p< 0.05 and ** p< 0.01 (comparisons as depicted in the Figure). Validation of fibronectin knockdown and ITGB1 over-expression is also shown. Data are representative of 3 independent experiments, n=3, Mean ± SEM.

### DDR2 knockdown attenuates Ang II-stimulated expression of AT1R in cardiac fibroblasts

Since fibronectin expression is dependent on DDR2, we investigated the effect of DDR2 knockdown on Ang II-stimulated AT1R expression. siRNA-mediated DDR2 silencing reduced both basal and Ang II-stimulated AT1R expression (Figure 5A). However, DDR2 over-expression failed to restore AT1R in fibronectin-silenced cells, showing that DDR2 regulates AT1R via fibronectin (Figure 5B).

**Figure 5:**
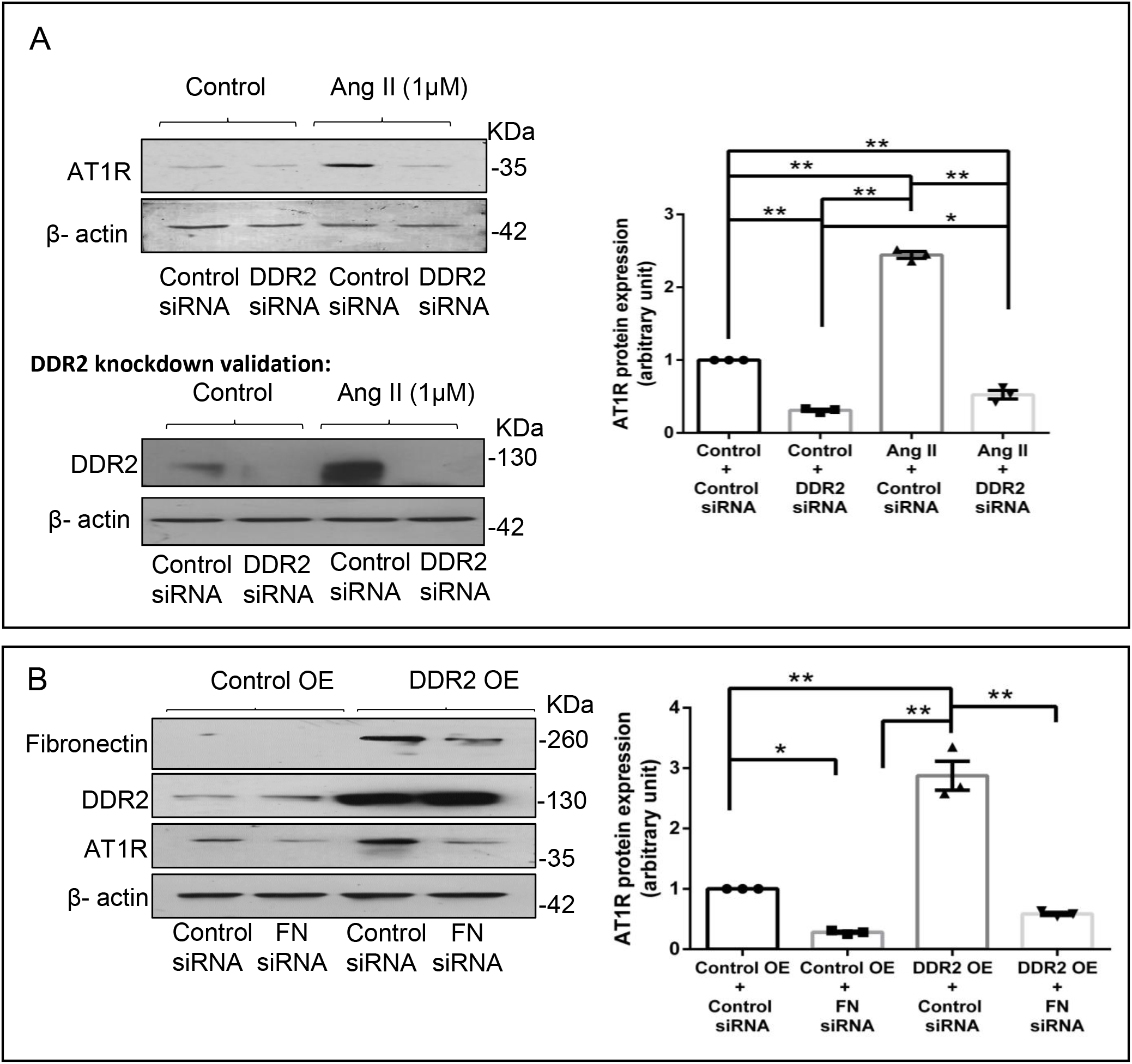
DDR2 regulates Ang II-stimulated expression of AT1R in cardiac fibroblasts. (**A**) Cardiac fibroblasts were transiently transfected with DDR2 siRNA (5 pmol) or control (scrambled) siRNA prior to treatment with Ang II for 12 h. AT1R protein expression was examined, with β-actin as loading control. Validation of DDR2 silencing is also shown. *p< 0.05, **p< 0.01 (comparisons as depicted in the Figure). (**B**) Cardiac fibroblasts were co-transfected with fibronectin siRNA (10 pmol) and DDR2 cDNA over-expression plasmid (2 μg). Following transfection, the cells were revived in M199 with 10% serum for 12 h. Post-revival, the cells were serum-deprived for 24 h prior to treatment with Ang II for 12 h. Cells were collected and AT1R protein levels were examined, with β-actin as loading control. *p< 0.05 and **p< 0.01 (comparisons as depicted in the Figure). Validation of fibronectin knockdown and DDR2 over-expression is also shown. Data are representative of 3 independent experiments, n=3, Mean ± SEM.

### AT1R levels in DDR2-null mouse heart

In view of the obligate regulatory role of DDR2 in Ang II-stimulated cardiac fibroblasts, we examined AT1R levels in DDR2-knockout mice carrying a germline deletion of DDR2. Knockin of a MerCreMer gene targeting exon 3 of the DDR2 allele was used for germ-line deletion of DDR2 in mice, and we had previously described the generation and validation of these mice, along with initial observations (10). In the present study, immunohistochemical analysis of cardiac tissue sections using an anti-AT1R antibody demonstrated a small but significant reduction in myocardial AT1R expression in DDR2-null mice (Figure 6A). However, the global reduction in AT1R staining intensity suggested a likely decrease in AT1R in myocytes in addition to fibroblasts, which are reportedly fewer in mouse heart (29).

**Figure 6:**
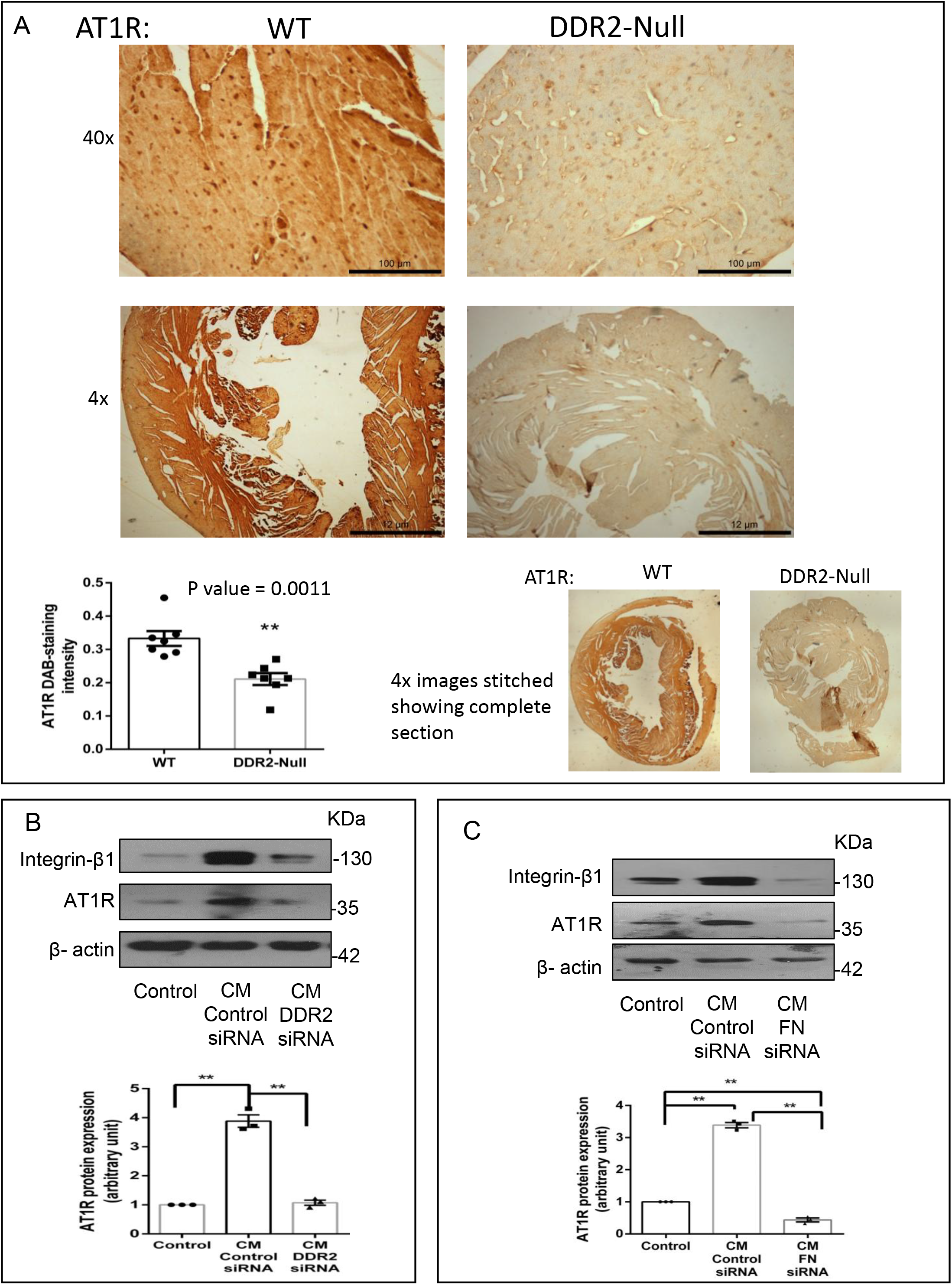
Corroboration of the DDR2-AT1R link *in vivo*. (**A**) Representative image showing 3,3’-diaminobenzidine (*DAB*) staining of AT1R protein in myocardial tissue sections of 10-week-old WT and DDR2-null mice. \*\**p*<0.01 *versus* WT (*n* = 7). (**B**) Sub-confluent quiescent cultures of H9c2 cells were treated for 24 h with conditioned medium (*CM*) derived from control siRNA-treated cardiac fibroblasts or DDR2-silenced cardiac fibroblasts. Quiescent cultures of H9c2 in M199 without serum were used as control for basal AT1R protein expression in these cells. AT1R protein expression in H9c2 cells was examined by western blot analysis and normalized to β-actin. \*\**p*<0.01 (comparisons as depicted in the Figure), (*n*=3, for the conditioned medium experiments, cardiac fibroblasts were from 3 isolations from 3 rats). (**C**) Sub-confluent quiescent cultures of H9c2 cells were treated for 24 h with conditioned medium (*CM*) derived from control siRNA-treated cardiac fibroblasts or fibronectin (FN)-silenced cardiac fibroblasts. Quiescent cultures of H9c2 in M199 without serum were used as control for basal AT1R protein expression in H9c2 cells. AT1R protein expression was examined by western blot analysis and normalized to β-actin. \*\**p*<0.01 (comparisons as depicted in the Figure), (*n*=3, for the conditioned medium experiments, cardiac fibroblasts were from 3 isolations from 3 rats).

### Cardiac fibroblasts influence AT1R expression in H9c2 cells via DDR2-dependent paracrine mechanisms

Preliminary experiments were therefore performed to explore the influence of fibroblast-specific DDR2 on myocyte AT1R expression. To this end, the effect of rat cardiac fibroblast-conditioned medium on AT1R expression in the rat ventricular H9c2 cell line (cardiomyoblasts) was analyzed. Exposure of H9c2 cells to fibroblast-conditioned medium for 24 h enhanced AT1R expression, but notably, conditioned medium from DDR2-silenced (Figure 6B) or fibronectin-silenced (Figure 6C) cardiac fibroblasts did not enhance AT1R expression in H9c2 cells, pointing to a DDR2/fibronectin-dependent paracrine effect of cardiac fibroblasts on AT1 expression in myocytes.

A schematic representation of the molecular events that link ECM and Ang II signaling pathways in cardiac fibroblasts is provided in Figure 7.

**Figure 7:**
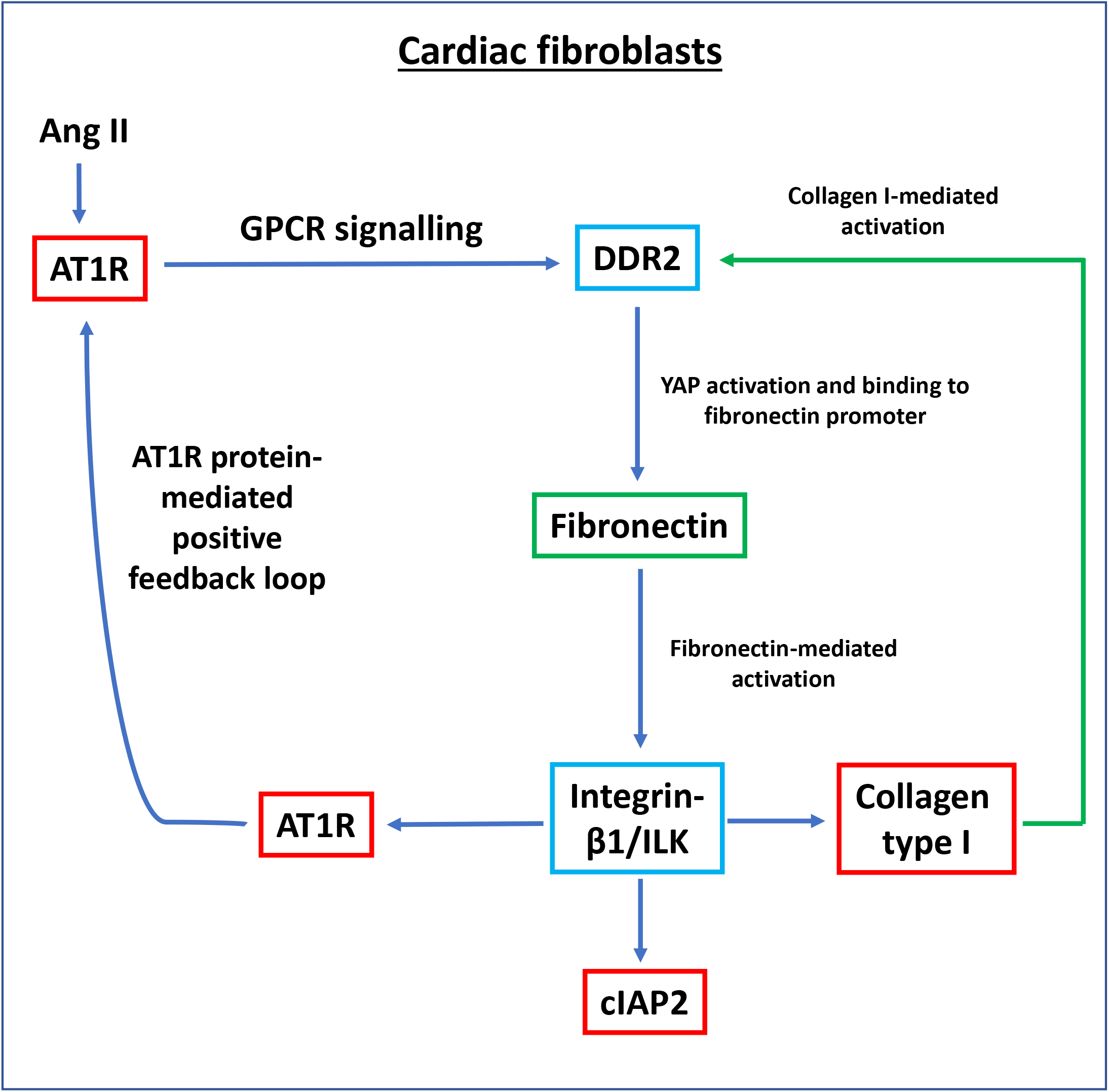
Schematic representation of the molecular events that link ECM and Ang II signaling pathways in cardiac fibroblasts.

## Discussion

Regardless of etiology, most myocardial diseases are associated with interstitial fibrosis due to excessive deposition of ECM proteins by cardiac fibroblasts and consequent impairment of the structural and functional integrity of the heart. Despite the centrality of interstitial fibrosis in heart disease and increasing appreciation that therapy directed at cardiac fibrosis may retard heart failure, there are limited therapeutic options to effectively target cardiac fibroblasts and attenuate cardiac fibrosis. Blocking the actions of Ang II either with AT1R blockers or ACE inhibitors is an accepted treatment strategy to reduce adverse cardiac remodelling following myocardial infarction but it may entail unfavourable effects such as the paradoxical activation of a negative feedback loop within the Renin-Angiotensin cascade, leading to elevation of systemic Ang II (30–32). Identification of alternate therapeutic targets would hinge around a sound understanding of the molecular basis of cardiac fibrogenesis.

In this regard, the critical role of ECM in myocardial pathophysiology, particularly in the pathogenesis of heart failure, has attracted increasing attention in recent years. ECM proteins significantly impact cellular responses, cell migration, differentiation, proliferation and survival through their ability to bind to multiple interacting partners, including other ECM proteins, and transmission of growth factor signals (33, 34). Alterations in ECM are reported to play an important role in modulating fibroblast phenotype and function (35). Major ECM proteins, collagen and fibronectin, and their receptors, DDR2 and integrins, participate in signaling events and determine cell fate (36). Our previous studies had shown that collagen type I-activated DDR2 is a ‘master switch’ in cardiac fibroblasts with an obligate role in phenotypic transformation, proliferation, collagen expression and apoptosis resistance in cardiac fibroblasts (2, 6, 7).

Fibronectin is a high molecular weight glycoprotein constituent of ECM that interacts with several matrix and cell surface proteins to regulate cell adhesion, migration, metastasis, proliferation and differentiation, as well as embryogenesis, wound healing, and blood coagulation (3). It has also been reported that tissue fibrosis is associated with up-regulation of fibronectin (16–19). In the present study, Ang II was found to enhance fibronectin expression (Figure 1A). Though Ang II per se has not been shown to enhance fibronectin, our observation is consistent with scleraxis-mediated enhancement of fibronectin gene expression in cardiac fibroblasts by TGF-β1 (37) since it is known that Ang II exerts some of its effects via TGF-β signaling in these cells (38).

We had shown previously that Ang II acts via the AT1 receptor and the GPCR pathway to enhance DDR2 expression that in turn plays an indispensable role in collagen type I gene expression (2). The findings reported here indicate that DDR2 mediates Ang II-stimulated expression of not only collagen type I but also of fibronectin (Figure 1B,C), which underscores its critical role in ECM homeostasis and ECM signaling. Further, our data show transcriptional up-regulation of fibronectin by DDR2-dependent activation of YAP (Figure 2), which is a mechano-sensitive transcriptional co-activator (39). The observation is consistent with previous studies that implicate transcriptional activation of YAP in pro-fibrotic gene expression in fibroblasts (39). Moreover, a previous study from our laboratory implicated Ang II-dependent activation of YAP in collagen type I expression (6), and inhibition of YAP has been reported to attenuate cardiac fibrosis (40).

It was of obvious interest to explore the pathophysiological significance of augmented fibronectin expression in cardiac fibroblasts. Cardiac fibroblasts are relatively resistant to several death signals that prevail in the injured myocardium, which enables these cells to play a central role in tissue repair following myocyte loss but also facilitates their persistence in the infarct scar, resulting in disproportionate stromal growth and pump dysfunction (1, 41). While a role for constitutively expressed Bcl-2 in the resistance of cardiac fibroblasts to a variety of pro-apoptotic stimuli was reported by Mayorga et al (42), our recent studies showed that anti-apoptotic cIAP2, induced by Ang II via a DDR2- and Serum Response Factor-mediated mechanism, protects cardiac fibroblasts from ambient stress (7). The present study extends these earlier observations and points to a role for fibronectin, downstream of DDR2, in Ang II-stimulated cIAP2 expression (Figure 3A) and protection of the cells from oxidative damage (Figure 3B). In a recent study by Valiente-Alandi et al, no significant change in fibroblast survival was observed upon inhibition of fibronectin polymerization under basal conditions (4). However, it is possible that fibronectin is important for survival under stress conditions.

Collagen type I is the most abundant ECM protein in the heart and is a major constituent of the scar tissue formed following injury (43). Excessive collagen deposition leads to diminished ventricular compliance that contributes to the initiation and progression of heart failure. Regulation of collagen expression has understandably been a subject of intense investigation for a long time. The findings presented here show that, apart from a role in cIAP2 expression, fibronectin has a role in collagen type I expression in Ang II-treated cells (Figure 3C), consistent with what was observed by Valiente-Alandi et al in an ischemia-reperfusion injury model (4).

Probing the regulatory role of fibronectin further, we found that its knockdown attenuates the expression of not only cIAP2 and collagen type I but also of Integrin-β1 (Figure 4A), which is known to be importantly involved in transducing ECM signaling (44). Thus, apart from acting as a ligand to trigger Integrin-β1 signaling, fibronectin also regulates Integrin-β1 gene expression in Ang II-stimulated cardiac fibroblasts. Further, plasmid-mediated over-expression of Integrin-β1 in cells following fibronectin knockdown restores, at least in part, the expression of cIAP2 and collagen type I (Figure 4D). Together, the data indicate that fibronectin is required for the expression of Integrin-β1 that in turn regulates cIAP2 and collagen type I in response to Ang II. The observed effect of Integrin-β1 over-expression in fibronectin-silenced cells suggests that fibronectin may not be indispensable as a ligand for Integrin-β1 signaling, and that other ligands such as collagen may also trigger Integrin-β1 signaling. Our earlier study had shown that a cross-talk between the two major collagen receptors, DDR2 and Integrin-β1, is a critical determinant of α-SMA-dependent collagen expression in Ang II-stimulated cardiac fibroblasts (6). Moreover, cell survival through ECM-integrin interactions has long been recognized (45). In light of the data presented here, it appears that fibronectin acts downstream of DDR2 to regulate Integrin-β1 signaling-dependent gene expression in cardiac fibroblasts. Our findings on the role of Integrin-β1 as an ‘effector molecule’ in a complex regulatory cascade are consistent with an earlier report that ECM-mediated mechanical stress activates TGF-β and triggers expression of Integrins and the pro-fibrotic gene program in cardiac fibroblasts (46).

We also examined the mechanistic basis of fibronectin-mediated apoptosis resistance and collagen expression in Ang II-stimulated cardiac fibroblasts, downstream of DDR2. It is known that Ang II, an effector molecule of the renin-angiotensin system, acts via the AT1 receptor to exert its potent pro-fibrotic effects on cardiac fibroblasts, which makes Ang II-AT1 signaling a major regulator of cardiac fibroblast response to myocardial injury, and hence a determinant of structural and functional alterations in the injured heart (47). We had previously uncovered a complex mechanism of regulation of AT1R involving the redox-sensitive NF-κB and AP-1 transcription factors that are activated by the co-ordinated action of ERK1/2 MAPK, p38 MAPK and JNK in cardiac fibroblasts exposed to oxidative stress (22). Interestingly, Ang II produced endogenously in response to oxidative response was found to be responsible for increased AT1R expression (22). Here, we present the novel finding that fibronectin has an obligate role in the regulation of AT1R in Ang II-stimulated cardiac fibroblasts (Figure 3D). Notably, while knockdown of Integrin-β1 or integrin-linked kinase attenuated Ang II-stimulated AT1R expression, over-expression of Integrin-β1 in fibronectin-silenced cells partially restored AT1R expression (Figure 4D), showing that Integrin-β1 signaling underlies, at least in part, the regulatory role of fibronectin in enhanced AT1R expression in Ang II-stimulated cardiac fibroblasts. Consistent with the fact that both fibronectin and Integrin-β1 are under the regulatory control of DDR2, Ang II-stimulated AT1R expression was found to be attenuated in DDR2-silenced cells (Figure 5A). However, DDR2 over-expression did not restore AT1R in fibronectin-silenced cells, showing that DDR2 regulates AT1R via fibronectin (Figure 5B).

The relationship between DDR2 and AT1R was further evident from the reduced AT1R levels in myocardial tissue from DDR2-null mice (Figure 6). Admittedly, the observed difference was modest, though significant, but it is pertinent to note that these were basal levels, and hence noteworthy. Nonetheless, the regulatory role of DDR2 in AT1R expression in Ang II-stimulated cells could be far more pronounced, as borne out by the findings presented here. Our attempts to determine fibronectin levels in fixed myocardial tissue were not successful, possibly due to antigen masking, and it is also likely that only the injured myocardium may provide robust signals. However, we had previously reported a significant reduction in collagen (10) and Integrin-β1 (6) in cardiac tissue from DDR2-null mice. Additionally, data from the conditioned medium experiments presented here show that a loss of DDR2 or fibronectin in cardiac fibroblasts could repress paracrine factors regulating AT1R expression in myocytes. Considered in tandem, these data demonstrate the importance of the DDR2-fibronectin-Integrin-β1 axis in AT1R expression, pointing to a convergence of ECM signaling and Ang II signaling, and unravels a novel mechanism by which ECM can impact cardiac fibroblast function via Ang II signaling.

Our findings indicate the existence of a complex mechanism of regulation of cardiac fibroblast function involving two major ECM proteins, collagen type I and fibronectin, and their respective receptors, DDR2 and Integrin-β1. Future investigations should evaluate the possible role of other factors like periostin within this complex mechanistic framework of molecular events under the regulatory control of DDR2 in cardiac fibroblasts.

Lastly, as noted earlier, targeting ECM proteins could offer alternate therapeutic strategies to prevent or minimise adverse cardiac remodelling in pathological states. In this regard, it is significant that fibronectin inhibition is reported to exert cardioprotective effects through a significant reduction in cardiac fibrosis and adverse fibrotic remodelling in an ischemia/reperfusion injury model (42). Viewed in tandem with these earlier observations in an in vivo model of myocardial injury, our findings support the postulation that increased susceptibility of cardiac fibroblasts to apoptosis and prevention of excessive collagen deposition may, in part, explain the cardioprotective effect of fibronectin inhibition observed in the animal model. Based on their findings, Valiente-Alandi et al suggest that targeting fibronectin polymerization may be a new therapeutic strategy for treating cardiac fibrosis and heart failure. However, given the ubiquitous distribution and critical role of fibronectin and Integrin-β1 in the heart, targeting them may entail global consequences, compromising cardiac integrity and function. On the contrary, since fibronectin and Integrin-β1 are under the regulatory control of DDR2, it is logical to postulate that the predominant localization and regulatory role of DDR2 in cardiac fibroblasts mark it as a cardiac fibroblast-specific drug target to control cardiac fibrosis.

## Experimental procedures

### Materials

Angiotensin II and M199 were obtained from Sigma-Aldrich, (St. Louis, MO, USA). Lipofectamine 2000 was from Invitrogen (Carlsbad, CA, USA). The Low Cell# ChIP kit protein A × 48 was from Diagenode (Denville, NJ, USA) and the Chemiluminescence western blot detection reagent was from Thermo Fisher Scientific (Waltham, MA, USA). DDR2, Integrin-β1, ILK and control siRNAs were from Ambion (Foster City, CA, USA). Fibronectin siRNA was custom-synthesized by Eurogentec (Liège, Belgium). The rat DDR2/CD167b Gene ORF cDNA clone expression plasmid was obtained from Sino Biologicals (Beijing, China). Pcax Itgb1-FLAG was a gift from Dennis Selkoe and Tracy Young-Pearse (Addgene plasmid 30153) (8). Opti-MEM and fetal calf serum (FCS) were from GIBCO (Waltham, MA, USA). All cell culture ware was purchased from BD Falcon (Corning, NY, USA). Primary antibodies against DDR2 (Cat No. 12133S), Caspase 3 (Cat No. 9662) and cleaved-caspase 3 (Cat No. 9661) were obtained from Cell Signaling Technology (Danvers, MA, USA). Primary antibodies against cIAP2 (Cat No. sc7944), Fibronectin (Cat No. sc9068), AT1R (Cat No. sc1173) and Collagen I (Cat No. sc293182) were from Santa Cruz Biotechnology (Dallas, TX, USA). Loading control β-Actin (Cat No. A2228) antibody was obtained from Sigma-Aldrich (St. Louis, MO, USA). ILK antibody was obtained from Elabscience (Houston, Texas, USA). All antibodies were used after dilution (1:1000). XBT X-ray Film was from Carestream (Rochester, NY, USA). The study on rats was approved by the Institutional Animal Ethics Committees of Sree Chitra Tirunal Institute for Medical Sciences and Technology, and the study on mice was approved by the Institutional Animal Care and Use Committee of the University of California, San Diego.

### Methods

#### Isolation of cardiac fibroblasts

Cardiac fibroblasts were isolated from young adult male Sprague–Dawley rats (2–3 months old) as described earlier (9). Sub-confluent cultures of cardiac fibroblasts from passage 2 or 3 were used for the experiments. Cells were serum-deprived for 24 h prior to treatment with 1 μM Ang II. Cells were pre-incubated with 10 μM Verteporfin for 1 h before the addition of 1 μM Ang II in the appropriate group. The cells were collected and processed further to analyse the expression of various genes and proteins.

#### Western blot analysis

Sub-confluent cultures of cardiac fibroblasts in serum-free M199 were treated with Ang II (1 μM) and relative protein abundance was determined by western blot analysis following standard protocols, with β-actin as the loading control. Enhanced chemiluminescence reagent was used to detect the proteins with X-ray Film.

#### RNA interference and over-expression

Cardiac fibroblasts at passage 3 were seeded on 60 mm dishes at equal density. After 24 h, the cells were incubated in Opti-MEM for 5–6 h with Ambion pre-designed Silencer-Select siRNA or custom-designed siRNA from Eurogentec or scrambled siRNA (control siRNA) at the given concentrations (10 pmoles for both DDR2 and Integrin-β1, 20 pmoles for Fibronectin (mix of siRNA 1 and 2) and Lipofectamine 2000.

Constitutive expression of DDR2 and Integrin-β1 was achieved under the control of a CMV promoter. Both plasmids were verified by restriction mapping. For over-expression, the plasmid vector for DDR2 (2 μg/μl) was transfected using Lipofectamine 2000. Following a post-transfection recovery phase in M199 with 10% FCS for 12 h, the cells were serum-deprived for 24 h and then treated with Ang II (1 μM) for the indicated durations. Cell lysates were prepared in Laemmli sample buffer, denatured and used for western blot analysis.

#### Chromatin Immunoprecipitation (ChIP) assay

The ChIP assay was performed with the Low Cell Number ChIP kit, according to the manufacturer’s protocol. Briefly, after treatment of cardiac fibroblasts with 1 μM Ang II for 30 min, the cells were cross-linked with 1% formaldehyde, lysed and sonicated in a Diagenode Bioruptor to generate ~600 bp DNA fragments. The lysates were incubated with anti-YAP antibody overnight at 4°C with rotation. Immune complexes were precipitated with protein A-coated magnetic beads. After digestion with proteinase K to remove the DNA-protein cross-links from the immune complexes, the DNA was isolated and subjected to PCR using primers for the specific promoter regions. In samples immunoprecipitated with the YAP antibody, the fibronectin promoter region was amplified using FP-5’-AAAACCGTTTTGTCAAGGGATG-3’ and RP-5’-TACCAGTTTCTTACAAGCGGTG-3’. DNA isolated from an aliquot of the total sheared chromatin was used as loading control for PCR (input control). ChIP with a non-specific antibody (normal rabbit IgG) served as negative control. The PCR products were subjected to electrophoresis on a 2% agarose gel.

#### In vivo study and histology

The generation of DDR2-null mice and genotyping were described in our previous study (10). Mouse tissue sections were prepared as described previously (10). Briefly, hearts were collected from age-matched (10 weeks) WT and DDR2-null mice, fixed in 4% buffered paraformaldehyde for 2 days, embedded in paraffin, cross-sectioned, and mounted onto slides. AT1R levels in these sections were analyzed by 3,3-diaminobenzidine staining and quantified using Fiji-ImageJ software.

#### Conditioned medium experiments

Rat ventricular H9c2 cells were used as an *in vitro* model for myocytes. The effect of cardiac fibroblast-conditioned medium on H9c2 cells was studied. Cardiac fibroblasts isolated from young adult (2-3 months) male Sprague Dawley rats were transfected with scrambled siRNA, DDR2 siRNA or fibronectin siRNA. Six hours post-transfection, the cells were revived in M199 with 10% serum for 12 h. Subsequently, the cells were transferred to M199 without serum for 24 h to generate fibroblast-conditioned medium. H9c2 cells were exposed to the fibroblast-derived conditioned medium from each of the transfected groups, collected after 24 h of incubation. H9c2 cells exposed to fresh M199 medium without serum served as controls for basal expression. Lysates of H9c2 cells were collected at 24 h following exposure to the conditioned medium, and AT1R protein expression was analyzed and quantified after normalization to β-actin expression.

#### Statistical analysis

Data are expressed as Mean ± SEM (Standard Error of Mean). Statistical analysis was performed using Student’s *t* test (unpaired, 2-tail) for comparisons involving 2 groups. For comparisons involving more than 2 groups, the data were analyzed by one-way ANOVA (with one variable) or two-way ANOVA (with two variables). p≤0.05 was considered significant. The in vitro data presented are representative of 3 or 4 independent experiments and in vivo data are representative of 7 age-matched animals from each group.

## Acknowledgements

This work was supported by a Research Grant (BT/PR23486/BRB/10/1589/2017) to SK from the Department of Biotechnology, Government of India, and Research Fellowships from SCTIMST, Trivandrum, to AST and the Department of Biotechnology, Government of India, to HV. The source of funding for MW and EGL was the Intramural Research Program of the National Institute on Aging, National Institutes of Health. SK, AST and HV thank Dr Ajay Kumar R of the Rajiv Gandhi Centre for Biotechnology, Trivandrum, for providing access to the Bioruptor Facility and acknowledge the facilities provided by SCTIMST, Trivandrum.

## Declaration

### Author contributions

AST, SK, EGL, HV and MW conceived the study; AST and SK designed the study; AST, HV and MGU performed the experiments; RTC generated the DDR2-null mice and prepared the heart tissue; AST, HV and SK analyzed the data; AST, HV and SK interpreted the results of the experiments; AST prepared the Figures; AST and SK drafted the manuscript; AST, HV, MGU, MW, EGL, RTC and SK reviewed the manuscript and approved the final version of the manuscript.

